# Metastatic breast cancers have reduced immune cell recruitment but harbor increased macrophages relative to their matched primary tumors

**DOI:** 10.1101/525071

**Authors:** Li Zhu, Jessica L. Narloch, Sayali Onkar, Marion Joy, Catherine Luedke, Allison Hall, Rim Kim, Katherine Pogue-Geile, Sarah Sammons, Naema Nayyar, Ugonma Chukwueke, Priscilla K. Brastianos, Carey K. Anders, Adam C. Soloff, Dario AA Vignali, George C. Tseng, Leisha A. Emens, Peter C. Lucas, Kimberly L. Blackwell, Steffi Oesterreich, Adrian V. Lee

## Abstract

The interplay between the immune system and tumor progression is well recognized. However, current human breast cancer immunophenotyping studies are mostly focused on primary tumors with metastatic breast cancer lesions remaining largely understudied. To address this gap, we examined exome-capture RNA sequencing data from 50 primary breast tumors (PBTs) and their patient-matched metastatic tumors (METs) in brain, ovary, bone and gastrointestinal tract. We used gene expression signatures as surrogates for tumor infiltrating lymphocytes (TIL) and compared TIL patterns in PBTs and METs. Enrichment analysis and deconvolution methods both revealed that METs have a significantly lower abundance of total immune cells, including CD8+ T cells, regulatory T cells and dendritic cells. An exception was M2-like macrophages, which were significantly higher in METs across the organ sites examined. Multiplex immunohistochemistry results were consistent with data from the in-silico analysis and showed increased macrophages in METs. We confirmed the finding of a significant reduction in immune cells in brain (BRM) METs by pathologic assessment of TILs in a set of 49 patient-matched pairs of PBT/BRMs. These findings indicate that METs have an overall lower infiltration of immune cells relative to their matched PBTs, possibly due to immune escape. RNAseq analysis suggests that the relative levels of M2-like macrophages are increased in METs, and their potential role in promoting breast cancer metastasis warrants further study.

## Introduction

Breast cancer is a highly heterogenous disease affecting 1 in 8 women in the US, and the most commonly diagnosed cancer in women worldwide. Despite recent improvements in overall survival rates, it is still the second leading cause of mortality due to cancer in women (1). In the last two decades, significant progress has been made in the detection and treatment of primary breast tumors as a result of enhanced understanding of disease biology and the tumor microenvironment (TME). The breast TME represents a complex interaction between tumor cells, endothelial cells, fibroblasts, and a variety of pro-and anti-tumor immune cells capable of tipping tumor biology toward tumor growth and progression or immune rejection. During tumor growth, cancer cells can be detected and eliminated by the immune system, but some cancer cells may exploit several mechanisms to evade destruction by the immune system, enabling them to escape immune surveillance and progress through the metastatic cascade. For breast cancer, the most common sites of distant organ metastases include bones, lungs, liver and brain with ovaries and gastrointestinal tract being affected less frequently (2).

The interplay between the immune system and tumor development is now well recognized in a variety of tumor types, including the triple negative (TNBC) and HER2+ subtypes of breast cancer (3,4). However, existing immunophenotyping studies focus mainly on primary tumors, with the role of immune cells in metastatic progression remaining largely understudied. While numerous studies have now documented cellular and genomic evolution of breast cancers during metastasis (5,6), very little is known about the co-evolution of immune cells and the TME. This study focused on addressing this gap in our understanding by performing immunophenotyping on two datasets: a) Pan-MET, transcriptomic profiles of 50 pairs of patient-matched primary (PBTs) and metastatic breast tumors (METs) in brain (BRM), ovary (OVM), bone (BOM) and gastrointestinal tract (GIM); and b) BRM-sTIL, a multi-institutional cohort of 49 patient-matched pairs of PBTs and BRMs with stromal tumor infiltrating lymphocytes (sTILs) percentages evaluated by pathologic evaluation of hematoxylin & eosin (H&E) staining. Using gene expression signatures as surrogates for TILs, we discovered quantitative differences in immune cell profiles between PBTs and METs in the first dataset (Pan-MET). Those differences were confirmed using multiplexed immunofluoresence (mIF) in three pairs of PBT/OVMs and PBTIBRMs each. Consistent results were observed by comparing the sTILs percentages in additional PBTIBRM pairs in a second dataset (BRM-sTIL). Higher immune cell recruitment to the TME was also confirmed to be associated with better survival in both datasets. Our study demonstrates the potential of using bioinformatics tools to investigate the evolution of the immune TME in breast cancer metastasis, and identifies M2-like macrophages as a potential therapeutic target for metastatic breast cancer.

## Materials and Methods

### Pan-MET dataset

Ethics approval and consent to participate were approved under the University of Pittsburgh IRE # PROIII00645, 14040193, 1500502, 1602030. Exome-capture RNA sequencing (ecRNA-seq) of patient-matched PBTs and METs collected from brain and bone were previously reported (7, 8). Ovarian and GI METs were recently reported (9). Clinical and pathological information of all samples are available in Supplementary Table S1. Formalin fixed paraffin embedded (FFPE) tissue sections of three pairs of PBT/BRMs and PBT/OVMs each were retrieved from the Pitt Biospecimen Core.

### BRM-sTIL dataset

Sample tissues of 49 pairs of patient-matched PBTs and BRMs were collected from four participating academic institutions (Duke University Medical Center, University of North Carolina Medical Center, University of Pittsburgh, Massachusetts General Hospital). Clinical and pathological information is available in Supplementary Table S2. 15 pairs of PBT/BRMs overlap between the Pan-MET and BRM-sTIL (Table S4).

### Immune abundance quantification of samples in Pan-MET dataset

Total immune single-sample gene set enrichment analysis (ssGSEA) score and tumor purity were calculated using R package ESTIMATE (10). Abundance of each immune cell population were calculated by R package GSVA (11) based on two sets of immune gene signatures, Davoli signatures (12) and Tamborero signatures (13). We also applied two deconvolution methods, CIBERSORT (14) and TIMER (15). All four methods were tested on a single cell RNA-seq dataset of 11 breast cancer tumors (16). If multiple METs were available for one patient, for a fair comparison of all pairs, average abundance was used for all comparisons.

### Differential expression (DE) test and pathway enrichment analysis

DE genes in ER+ BRMs versus PBTs were tested using R package DESeq2 (17). Significantly up-or down-regulated genes were further used for pathway enrichment analyses. We obtained 2531 pathways, contributed by BioCarta, GO, KEGG, Reactome, containing 5-2000 genes, from Molecular Signature Database (MSigDB Version 5.1. Broad Institute, Cambridge, MA, USA). Fisher’s exact test was performed with false discovery rate (FDR) 0.05 as cutoff.

### Multiplex staining experiment of selective pairs in Pan-MET dataset

FFPE tissue sections (5micron) were mounted on slides and deparaffinized. Briefly, tissues were subjected to cycles of antigen retrieval, blocking, primary antibody followed by secondary-HRP antibody. Separate Opal detection and signal amplification antibodies were used for each marker. The panel of markers used included CD8, CD20, CD68, Foxp3, PD-L1, pan-CK and DAPI. Imaging, analysis and quantification was performed using the Perkin Elmer Vectra platform and Inform software (18). The list of antibodies with catalog numbers and dilutions used provided in supplementary Table S5.

### Evaluation of stromal tumor infiltrating lymphocytes (sTILs) in BRM-sTIL dataset samples

H&E stained sections were manually counted for percent sTILs using standard criteria developed by the international TILs working group (19). sTILs were rounded to the nearest 5% increment. Only the stromal compartment within the borders of invasive tumor was evaluated. TILs in zones of necrosis, crushed artifacts, or normal tissue were excluded. Only mononuclear infiltrate was counted. Full tumor sections were preferentially examined over needle biopsies whenever possible; core biopsies were analyzed when full sections were unavailable. Each slide was independently reviewed by two study personnel (JLN and CL) to minimize inter-observer variability. When the sTILs differed by 10% or more, the study pathologist (AR) made the final determination. If multiple BRMs or PBTs were available for the same patient, average sTILs percentage was used for all comparisons.

### Data and software availability

Code and data for all bioinformatic analyses are available on https://github.com/lizhu06/TILsComparison_PBTvsMET.

## Results

### METs have lower total immune abundance than patient-matchedPETs

We estimated total immune abundance using RNAseq from 50 pairs of patient-matched PBTs and METs. METs showed a significantly lower total immune ssGSEA score compared to patient-matched PBTs (Fig 1A; p<0.001). To minimize the potential bias of tumor cellularity, we also compared the normalized ssGSEA immune scores which were divided by the percentage of non-tumor content, as higher tumor purity naturally indicates lower immune abundance. Since such normalization did not change the conclusion (Fig S1A), and considering this normalization was an overly simplified approximation and did not differentiate tumor from stromal regions, we used the immune ssGSEA score without normalization for further analysis to avoid introducing additional bias. The decrease in immune score was observed in METs collected from various sites, but was especially apparent in BRMs (Fig 1B). Validating this finding, pathologic assessment of sTILs in an an additional cohort of 49 patient-matched PBTs and METs revealed that BRMs also showed a significant decrease in the percentage of sTILs compared to patient-matched PBTs (Fig 1C; p<O.OOI). When grouping PBT/MET pairs by hormone receptor status (RR) and HER2 status, both datasets revealed a trend of decreased immune abundance in all subtypes, with TNBC subtype having the most significant decrease (Fig S1(B-C)). While the total immune ssGSEA score only estimates the overall immune abundance in the bulk sample from RNAseq, and the sTILs percentage was carefully counted as the immune cell percentage in the stroma, the two measurements of immune abundance were significantly (p< 0.001) correlated for the 15 pairs of PBTIBRMs within both data sets (Fig 1D).

**Figure 1.**
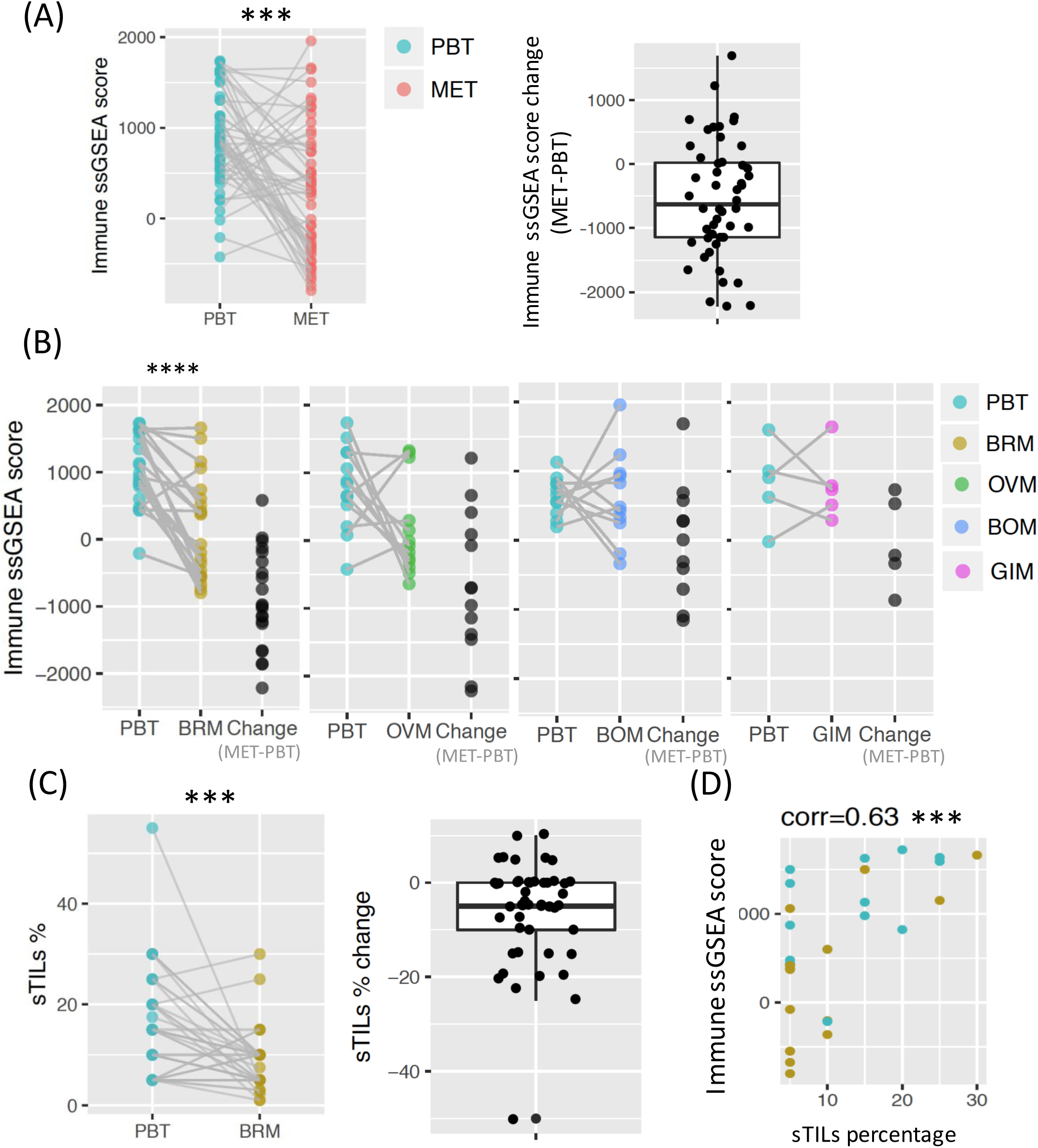
Lower immune abundance in metastatic breast tumors (METs) compared to primary breast tumors (PBTs). (A) Total immune ssGSEA score in PBTIMET pairs in Pan-MET dataset, together with the paired changes (MET-PBT). (B) Total immune ssGSEA score grouped by MET sites: METs in brain (BRM), ovary (OVM), bone (BOM), and GI (GIM). (C) Stromal tumor infiltrating lymphocytes (sTILs) percentages of 49 pairs of PBT/BRMs in BRM-sTIL dataset. (D) Spearman’s correlation between sTILs percentages and total immune ssGSEA score for 15 pairs of PBT/BRMs overlapped by Pan-MET and BRM-sTIL. ****p <0.0001, ***p <0.001, **p < 0.01, *p < 0.05 from two-sided Wilcoxon signed rank test in (A-C) and correlation test in (D)

In addition, we also observed that METs had significantly lower expression of immune checkpoint molecules that downregulate immune response-including CD274 (PD-Ll), PDCD 1 (PD-1) and CTLA4 (Fig S2)-possibly due to fewer total immune cells. Differential expression (DE) test identified 1,659 up-regulated and 1,036 down-regulated genes in the comparsion of ER+ BRMs versus PBTs under FDR 0.05 cutoff. Pathway enrichment analysis of DE genes identified KEGG pathway “primary immunodeficiency” among the top 15 significantly enriched pathways, further confirming our previous findings (Table S3).

Taken together, both transcriptomic data and pathological assessment showed that METs have lower immune abundance than patient-matched PBTs.

### METs have higher percentage of M2-like macrophages relative to the total immune abundance

We inferred the abundance of each immune cell population by enrichment analysis and deconvolution methods. To validate those approaches, we first compared the GSVA scores of four common immune cell populations defined by both Davoli et al. (12) and Tamborero et al. (13). The correlations ranged from 0.4 to 0.85 (Fig S3), indicating overall high consistency. For further validation, we applied four methods; namely GSVA using the immune signatures from Davoli and Tomborero, and two methods of deconvolution (CIBERSORT and TIMER) to a publicly available single cell RNA-seq dataset (16), in which immune cell percentages were available using cell markers. Based on the correlations, the estimated levels of B cell, T cell, and macrophages by immune signatures from Davoli and Tamborero, and deconvolution method TIMER, were in general most highly correlated with actual abundance of corresponding cell types, although some signatures were not quite specific, such as CD4+ mature T cell and CD8+ effector T cell in Davoli signatures. CIBERSORT estimates showed lower correlations as expected, because the actual percentages were calculated based on three cell types, while CIBERSORT considered 22 cell types (Fig S4).

Comparing patient-matched PBTs and METs, the GSVA score and abundance estimate from deconvolution methods for most immune cell populations were significantly lower in METs (Fig 2 A-C). Adjusting for total immune abundance, most immune cell populations were still lower, but M2-like macrophages were significantly higher in METs (Fig 2D). Significant increment was also observed in the ratio of the relative percentages of M2 and M1, indicating dominant level of M2 over M1 (Fig 2E). When separating PBTIMET pairs to different MET sites or HRlHER2 subtypes, the results were generally consistent (Fig S5 and S6).

**Figure 2.**
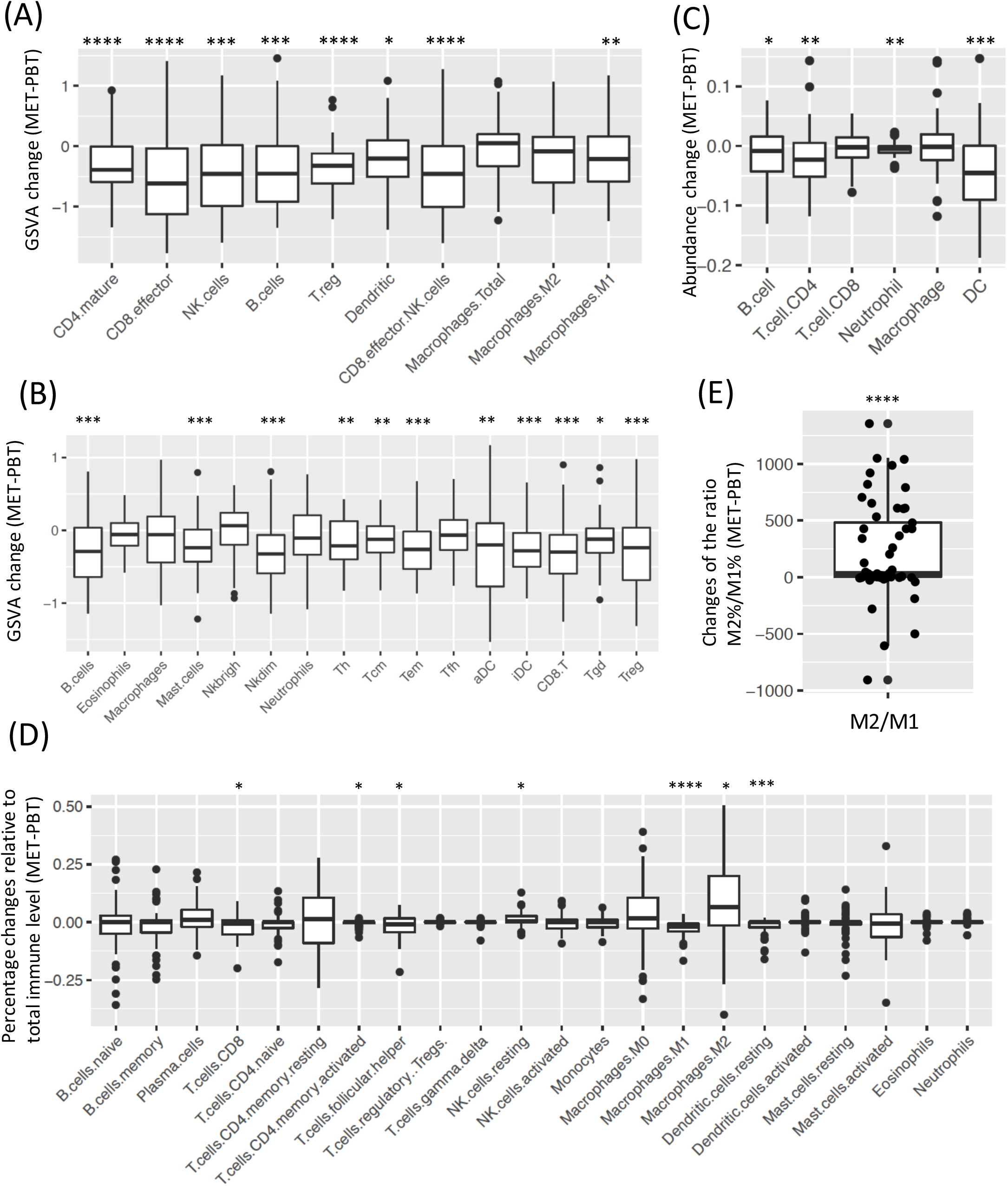
Paired comparison of the abundance of immune cell population in PBT/MET pairs in Pan-MET. (A-B) GSVA score changes (MET-PBT) of (A) Davoli signature and (B) Tamborero signature. (C) Abundance changes estimated by deconvolution method TIMER. (D) Changes of percentages relative to total immune level estimated by deconvolution method CIBERSORT. (E) Changes of the ratio of relative percentages of M2 and M1. ****FDR < 0.0001, ***FDR < 0.001, **FDR < 0.01, *FDR < 0.05 by Benjamini-Rochberg correction. Two-sided Wilcoxon signed rank test.

### Multiplex staining confirms the in-silica results

To further validate *in silico* results, we selected three pairs of PBTIBRMs and three pairs of *PBT/OVMs* which were shown to have higher M2-like macrophages relative to the total immune abundance. Multispectral immunofluorescence (mIF) was performed for the pairs (Fig 3A). Three pairs *of PBT/OVMs* and two pairs of PBTIBRMs showed increased macrophages in METs, and the majority of METs had lower B cells and T cells (Fig 3B), consistent with percentage estimated from CIBERSORT (Fig 3C and Fig S7).

**Figure 3.**
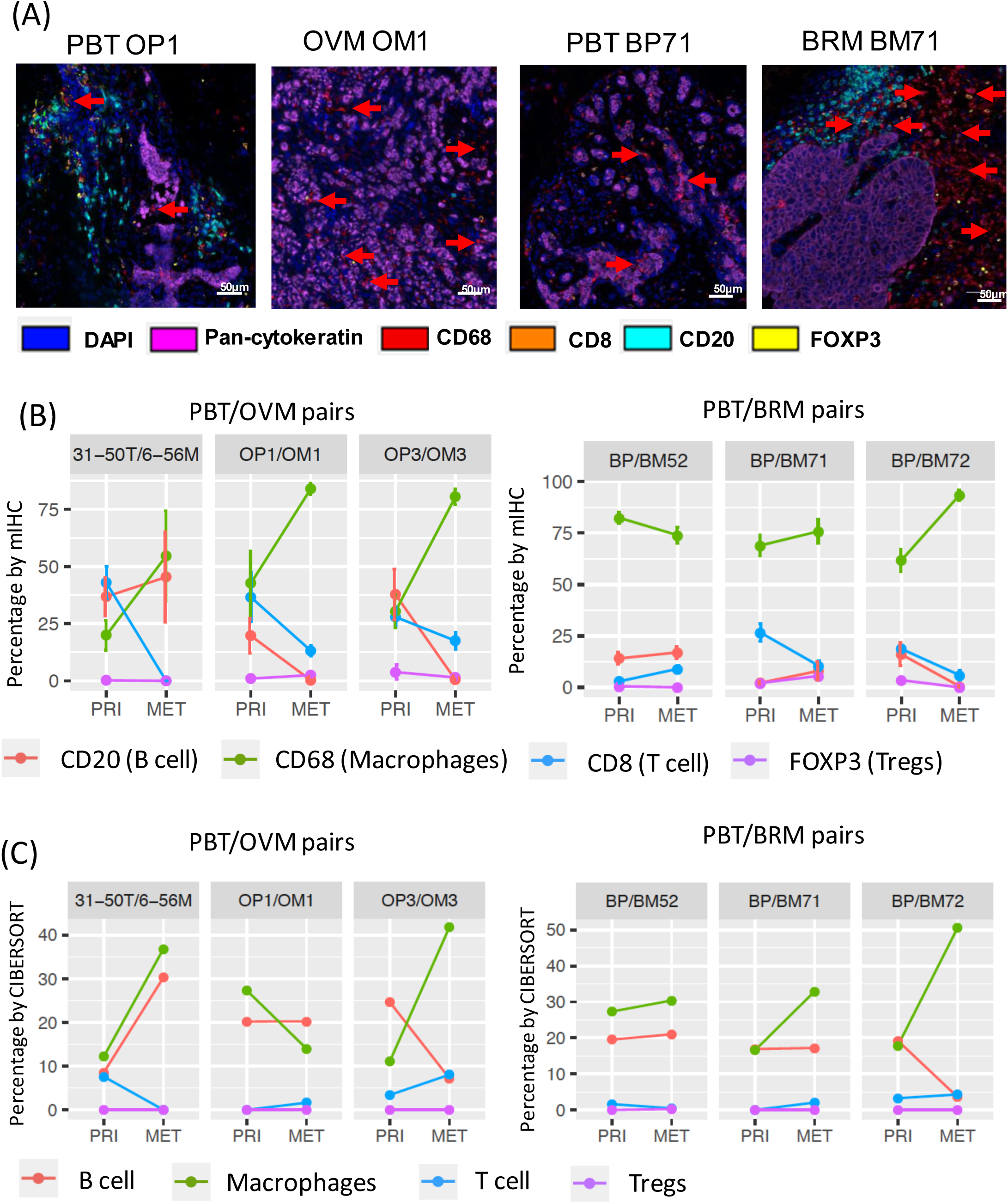
Multispectral immunohistochemical (mIRe) staining of selective pairs in Pan-MET. (A) mIRC staining images of one pair of PBT/OVMs and PBT/BRMs. (B) Percentage of each immune cell population denoted by markers using mIRC staining. (C) Relative percentages of corresponding immune cell populations estimated by CIBERSORT.

### Hormone receptor (HR) positive tumors are associated with lower total immune abundance

When examining associations between immune ssGSEA immune score and clinical variables, both the RNAseq and the sTIL dataset revealed that HR+ PBTs have significantly lower immune scores than HR-PBTs (Fig 4A). Further, HR+ METs tended to have a smaller decrease in immune abundance compared to PBTs, although this was only significant in the BRM-sTIL dataset.

**Figure 4.**
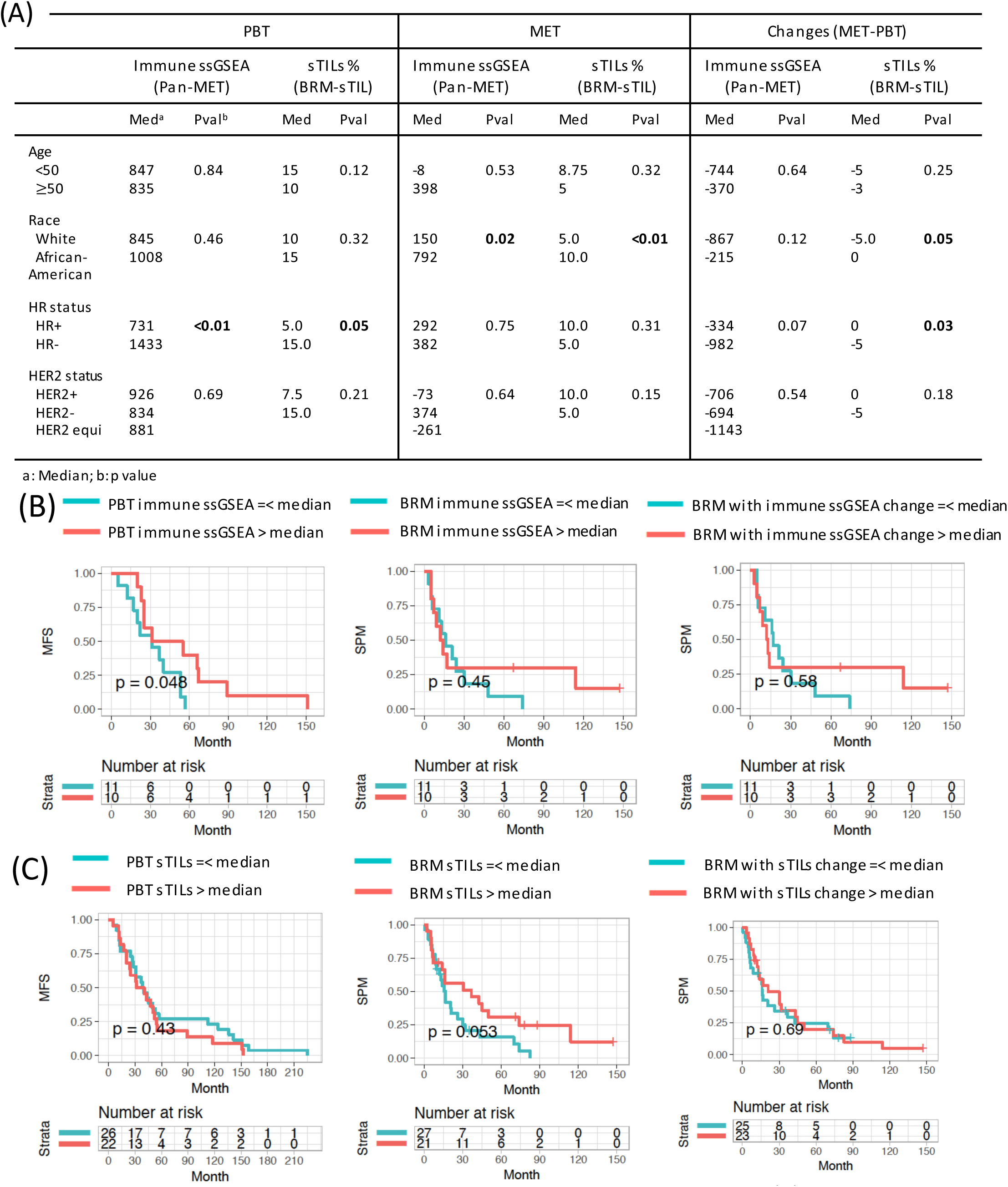
Association of immune abundance with clinical variables and survivals. (A) Association between immune ssGSEA score and sTILs with clinical variables. (B) Test association with survivals in Pan-MET datset: immune ssGSEA of PBTs and MFS, immune ssGSEA of BRMs and SPM, and immune ssGSEA changes and SPM. (C) Test associations with survivals in BRM-sTIL datset: sTILs of PBTs and MFS, sTILs of BRMs and SPM, and sTIL changes and SPM. ****p <0.0001, ***p <0.001, **p < 0.01, *p < 0.05 from Wilcoxon signed rank and Kruskal-Wallis test in (A) and log-rank test in (B)-(C).

### Higher immune abundance is associated with longer metastasis-free survival (MFS) and survival-post-metastasis (SPM)

We hypothesized that immune level of PBT may be associated with MFS, while immune level of MET and its change from PBT to MET are potentially associated with SPM. Combining all PBTIMET pairs into one cohort, immune ssGSEA score was not significantly associated with MFS or SPM (Fig S8), likely due to the confounding effect of different MET sites on outcome. Considering *PBTIBRM* pairs had the largest sample size, we tested the potential association between immune ssGSEA score and survival specifically in PBT/BRMs. In the pan-MET dataset, there was a trend in association between higher immune levels in PBTs and longer time to development of BRMs (i.e. MFS) (Fig 4B). However, such a trend was not observed between SPM with immune levels in BRM or immune level change between PBT and BRM (Fig 4B). In the BRM-sTIL dataset, higher sTILs percentage in PBT was not associated with MFS. Instead, there was a trend toward an association between a higher sTILs percentage in MET and longer SPM (Fig 4C). We did not observe significant associations between the relative level of M2-like macrophage and survivals (Fig S9).

## Discussion

It is now well appreciated that immune cells are a critical component of the TME. Studies of the breast TME have largely focused on tumor mutational and transcriptional landscapes in primary breast cancers, and with more recent attention to metastatic tumors. Our study is novel in two main regards: (1) we examined two cohorts of matched PBTs and METs, one of which includes METs in different sites, allowing us to discern site-specific immune changes from primary to metastatic disease and (2) we evaluated immune abundance by both gene expression analysis and H&E staining, and observed overall high consistency. Our data demonstrate the potential of using bioinformatics tools to investigate the immune contexture of both primary and matched metastatic tumors when tumor lesions may not be available for staining.

Our paired patient-matched comparison revealed a decrease in immune cells from primary to metastatic breast cancer, which is consistent with limited existing studies (20-22). In-silico analysis of the Pan-MET dataset, validated by mIF staining, highlights the potential enrichment of M2-like macrophages as the tumor cells metastasize to various sites, especially brain and ovary. This is consistent with the growing body of literature that has shown macrophages to be one of the key players in establishment of distant METs (23-25). Our survival analysis suggests enhanced MFS and SPM in patients with higher immune cell recruitment to primary and metastatic tumors, although the significance of these findings were not consistent between the Pan-MET and BRM-sTIL, possibly due to small sample size and/or sample heterogeneity.

This work has multiple important strengths. First, it utilizes established genomic data sets for elucidating the immunobiology of matched PBTs and METs. Second, it is one of the larger studies of a cohort of patient-matched PBTs and METs. Third, it effectively integrates state-of-the-art genomic analyses with multiplexed immunohistochemistry conducted in a subset of tumors to confirm results. Our study also has several limitations. First, due to the scarcity of patient-matched pairs of primary and metastatic breast cancer, our sample set remains somewhat small relative to studies of primary breast tumors alone. Second, RNAseq analysis was performed on bulk tumor samples, and thus gene expression cannot be attributed to specific cells. Although we attempted to reduce such bias by normalizing the immune ssGSEA score against the non-tumor cell percentage (with consistent conclusions), single cell RNA-sequencing may be needed to completely resolve uncertainties related to cellular heterogeneity. Third, in our mIF studies, the percentage of all immune cells within the tumor was often below 10%. Given these limited numbers of immune cells, our results should be interpreted with caution. Despite these limitations, our study clearly highlights an opportunity to utilize existing data to shed light on the co-evolution and involvement of immune cells in the progression of a primary tumor and its metastatic cascade within an individual patient. It also nominates M2-like macrophages as a potential target for therapeutic immune manipulation of the metastatic cascade.

## Acknowledgements

This project used the Pitt Biospecimen Core, and the UPMC Hillman Cancer Center Tissue and Research Pathology shared resource which is supported in part by NIHINCI award P30CA047904. We acknowledge the support of the UPMC Cancer Registry and in particular Louise Mazur for clinical data abstraction.

## Supplementary materials

**Table Sl. Clinical information of samples in Pan-MET dataset**

**Table S2. Clinical information of samples in BRM-sTIL dataset**

**Table S3. PathwayRes_MET_vs_PRI_Brain_ERpos.csv**

Pathway enrichment analysis of differentially expressed genes in comparison of ER+ Brain METs versus matched PBTs.

**Table S4. 15 pairs of PBT/BRMs overlap between the Pan-MET and BRM-sTIL.**

**Table S5. Detailed list of antibodies and dilutions used for multispectral immunofluorescence staining of slides as shown in Figure 3**

**Figure S1. Lower immune abundance in metastatic breast tumors (METs) compared to primary breast tumors (PBTs).**

(A) Comparison of normalized immune ssGSEA score in Pan-MET pairs. (B) Total Immune ssGSEA score in Pan-MET dataset, together with the paired changes (MET-PBT), grouped by HRlHER2 subtypes. (C) sTILs percentages of PBTIBRM pairs in BRM-sTIL dataset, grouped by HRlHER2 subtypes. ****p <0.0001, ***p <0.001, **p < 0.01, *p < 0.05 from two-sided Wilcoxon signed rank test.

**Figure S2. Expression (log2(TPM+1)) of CD274 (PD-L1), PDCD1 (PD-1), and CTAL4 in PBT and MET.**

Two-sided Wilcoxon signed rank test was used to compare PBT and MET. Spearman’s correlation with immune ssGSEA change was calculated and tested using correlation test. ****p <0.0001, ***p <0.001, **p < 0.01, *p < 0.05.

**Figure S3. Correlation between GSVA scores of Davoli and Tamborero signatures for PBT/MET pairs in Pan-MET dataset.**

**Figure S4. Correlation between immune abundance estimated from RNA-seq data and cell count/proportion (relative to total immune cell count) in single cell RNA-seq dataset.** (A-B) GSVA score of (A) Davoli and (B) Tamborero signatures. (C) Percentage relative to total immune level estimated by CIBERSORT. (D) Immune abundance estimated by TIMER. White in the heatmap indicates CIBERSORT estimates are all zero, and spearman’s correlation is not applicable.

**Figure S5. Comparison of the abundance of immune cell population in PBT/MET pairs in Pan-MET dataset, grouped by MET sites.**

(A-B) GSVA score change (MET-PBT) of (A) Davoli and (B) Tamborero signatures. (C) Abundance change estimated by deconvolution method TIMER. (D) Change of percentage relative to total immune estimated by deconvolution method CIBERSORT. ****FDR<O.OOOI, ***FDR <0.001, **FDR < 0.01, *FDR < 0.05. Two-sided Wilcoxon signed rank test.

**Figure S6. Comparison of the abundance of immune cell population in PBT/BRM pairs in Pan-MET dataset, grouped by HR/HER2.**

(A-B) GSVA score change (BRM-PBT) of (A) Davoli and (B) Tamborero signatures. (C) Abundance change estimated by deconvolution method TIMER. (D) Change of percentage relative to total immune estimated by deconvolution method CIBERSORT. ****FDR<O.OOOI, ***FDR <0.001, **FDR < 0.01, *FDR < 0.05. Two-sided Wilcoxon signed rank test.

**Figure S7. Correlation between mIHC staining results and CIBERSORT estimates.**

(A) *PBT/OVM* pairs. (B) *PBTIBRM* pairs in Pan-MET. Spearman’s correlation.

**Figure S8. Test association between survivals and total immune ssGSEA of all pairs of PBTs and METs in Pan-MET dataset.**

(A) Kaplan-Meier (KM) curves of MFS for PBTs with total immune ssGSEA below or above median. (B) KM curves of SPM for METs with total immune ssGSEA below or above median. (C) KM curves of SPM for METs with total immune ssGSEA change below or above median. P-values were from log-rank test.

**Figure S9. Test association between survivals and relative percentage of M2-like macrophages of PBT/BRM pairs in Pan-MET dataset.**

(A) Kaplan-Meier (KM) curves of MFS for PBTs with relative percentage of M2-like macrophage below or above median. (B) KM curves of SPM for METs with relative percentage of M2-like macrophage below or above median. (C) KM curves of SPM for METs with relative percentage change of M2-like macrophage below or above median. P-values were from log-rank test.

