## Supplemental figures and tables for "Metastatic breast cancers have reduced immune cell recruitment but harbor increased macrophages relative to their matched primary tumors"

### Supplementary tables and figures

### Table S1. Clinical information of samples in Pan-MET dataset

| Pairs | PBT/BRM (N=21) | PBT/OVM (N=12) | PBT/BOM (N=11) | PBT/GIM (N=5) | Total (N=49) |
|---|---|---|---|---|---|
| **Age (median (Q1[a],Q3[b]))** | 53 (40, 60) | 41 (38, 50) | 54(47, 58) | 51 (48, 57) | 50 (40, 57) |
| **Race (n, %[c])** | | | | | |
| **White** | 17 (81%) | 11 (100%) | 10 (91%) | 3 (60%) | 41 (85%) |
| **Black** | 4 (19%) | 0<br>1 missing | 1 (9%) | 2 (40%) | 7 (15%)<br>1 missing |
| **Post menopausal (n, %)** | 6 (55%)<br>10 missing | 3 (25%) | 5 (50%)<br>1 missing | 1 (25%)<br>1 missing | 15 (41%)<br>12 missing |
| **HR+ (ER+ or PR+) (n, %)** | 10 (48%) | 10 (83%) | 11 (100%) | 4 (80%) | 35 (71%) |
| **HER2+ (n, %)** | 8 (38%) | 1 (9%)<br>1 missing | 1 (9%) | 1 (20%) | 11 (23%)<br>2 missing |
| **Triple negative (n, %)** | 8 (38%) | 1 (10%)<br>2 missing | 0 | 1 (20%) | 10 (21%)<br>1 missing |
| **Histology (n, %)** | | | | | |
| **IDC** | 19 (90%) | 4 (33%) | 8 (73%) | 3 (60%) | 34 (69%) |
| **ILC** | 1 (5%) | 6 (50%) | 1 (9%) | 2 (40%) | 10 (20%) |
| **mixed** | 1 (5%) | 2 (17%) | 2 (18%) | | 5 (10%) |
| **Stage (n, %)** | | | | | |
| **I /II** | 12 (57%) | 10 (83%) | 4 (36%) | 2 (40%) | 28 (57%) |
| **III** | 6 (29%) | 2 (17%) | 4 (36%) | 3 (60%) | 15 (31%) |
| **IV** | 3 (14%) | 0 | 3 (27%) | 0 | 6 (12%) |
| **MFS (month median (Q1, Q3))** | 31 (22, 55) | 60 (21, 90)<br>1 missing | 24 (9, 40) | 60 (48, 69)<br>1 missing | 37 (20, 60)<br>2 missing |
| **SPM (month median (Q1, Q3))** | 14 (7, 30) | 29 (9, 57)<br>2 missing | 18 (3, 21) | 21 (19, 28)<br>1 missing | 18 (6, 39)<br>3 missing |

a: 25[TH] percentile; b: 75[th] percentile; c: Missing value were removed when calculating proportion;
c: MFS – metastasis free survival, time from primary diagnosis to recurrence of distant metastasis;
d: SPM – survival post metastasis, time from metastasis to death or last follow-up.

### Table S2. Clinical information of samples in BRM-sTIL dataset

| Pairs | PBT/BRM (N=49) |
|---|---|
| **Age (median (Q1[a],Q3[b]))** | 48 (39, 56) |
| **Race (n, %[c])** | |
| **White** | 31 (65%) |
| **African-American** | 13 (27%) |
| **Other** | 1 (2%) |
| **Unknown** | 3 (6%) |
| **HR+ (n, %)** | 25 (51%) |
| **HER2+ (n, %)** | 20 (47%) |
| **Triple negative (n, %)** | 15 (35%) |
| **Histology (n, %)** | |
| **IDC** | 47 (98%) |
| **ILC** | 1 (2%) |
| **MFS (in month, median (Q1, Q3))** | 40 (20, 56) |
| **SPM (in month, median (Q1, Q3))** | 16 (8, 43) |

a: 25$^{TH}$ percentile; b: 75$^{th}$ percentile; c: Missing value were removed when calculating proportion.

Table S4. 15 pairs of PBT/BRMs overlap between the Pan-MET and BRM-sTIL.

| ID | Site |
| --- | --- |
| BP12 | Breast |
| BM12 | Brain |
| BP17 | Breast |
| BM17 | Brain |
| BP19-2 | Breast |
| BM19-2 | Brain |
| BP25 | Breast |
| BM25 | Brain |
| BP29 | Breast |
| BM29 | Brain |
| BP47 | Breast |
| BM47 | Brain |
| BP51 | Breast |
| BM51 | Brain |
| BP52 | Breast |
| BM52 | Brain |
| BP6 | Breast |
| BM6 | Brain |
| BP62 | Breast |
| BM62 | Brain |
| BP64 | Breast |
| BM64 | Brain |
| BP68 | Breast |
| BM68 | Brain |
| BP7 | Breast |
| BM7 | Brain |
| BP71 | Breast |
| BM71 | Brain |
| BP72 | Breast |
| BM72 | Brain |

Table S5. Detailed list of antibodies and dilutions used for multispectral immunofluorescence staining of slides as shown in Figure 3

| | Reagents/ Antibodies used | Company | Cat# | Dilution used |
|---|---|---|---|---|
| 1 | Perkin Elmer 7 color manual Kit- including opal fluorophores and DAPI | Perkin Elmer | NEL811001KT | |
| 2 | CD8 - CLONE: C8/144B | Biocare Medical | ACI3160A | (1:200) |
| 3 | CD20 - CLONE: L-26 | Leica Biosystems | CD20-L26-L-CE | (1:200) |
| 4 | CD68 - CLONE: D4B9C | Cell Signaling | 76437S | (1:800) |
| 5 | Foxp3 - CLONE: D608R | Cell Signaling | 12653S | (1:250) |
| 6 | Pan-Cytokeratin - CLONE: AE1/AE3 | Santa Cruz Biotech | SC81714 | (1:100) |

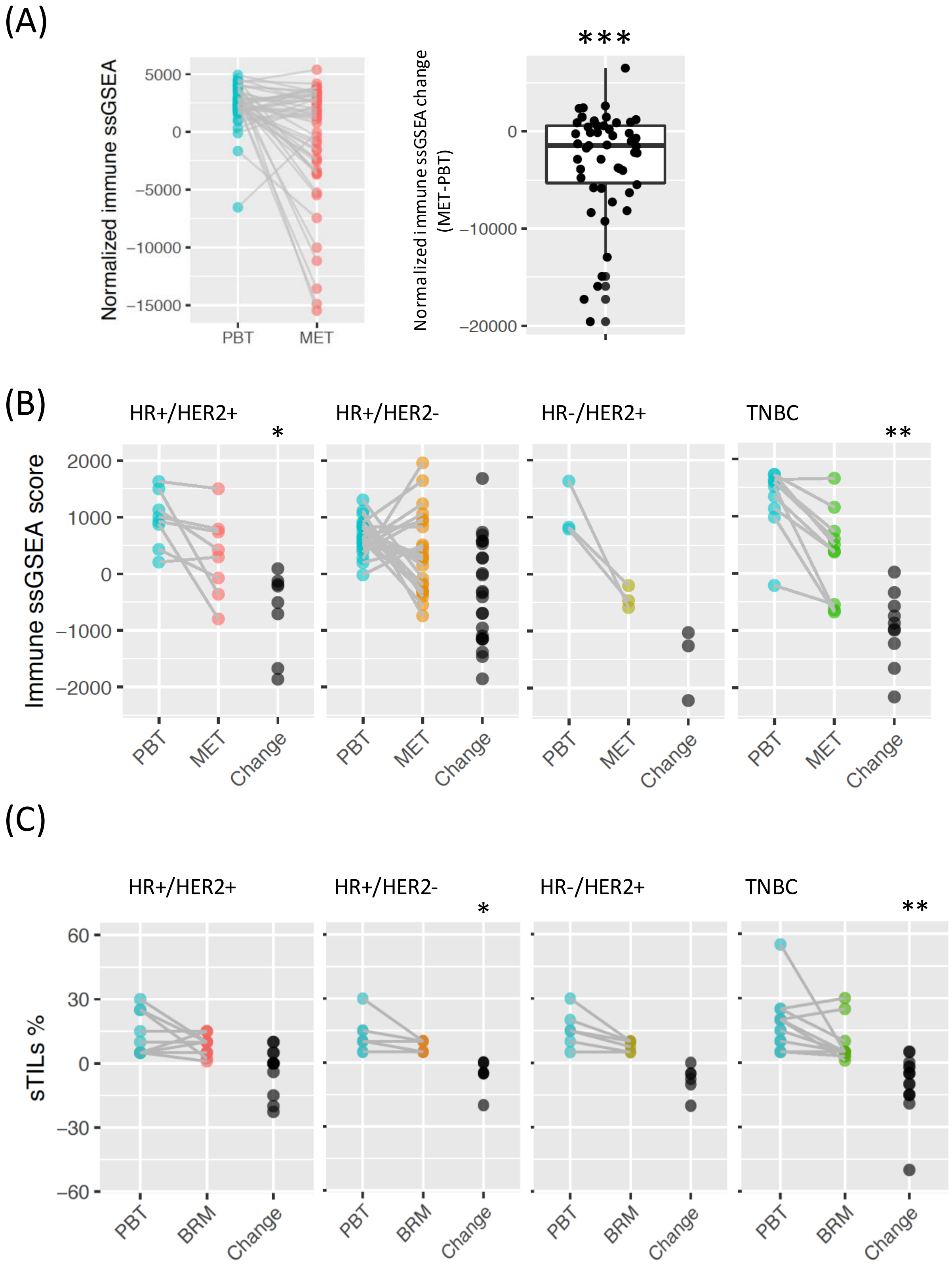


Figure S1. **Lower immune abundance in metastatic breast tumors (METs) compared to primary breast tumors (PBTs)**. (A) Comparison of normalized immune ssGSEA score in Pan-MET pairs. (B) Total Immune ssGSEA score in Pan-MET dataset, together with the paired changes (MET-PBT), grouped by HR/HER2 subtypes. (C) sTILs percentages of PBT/BRM pairs in BRM-sTIL dataset, grouped by HR/HER2 subtypes. ****p <0.0001, ***p <0.001, **p < 0.01, *p < 0.05 from two-sided Wilcoxon signed rank test.

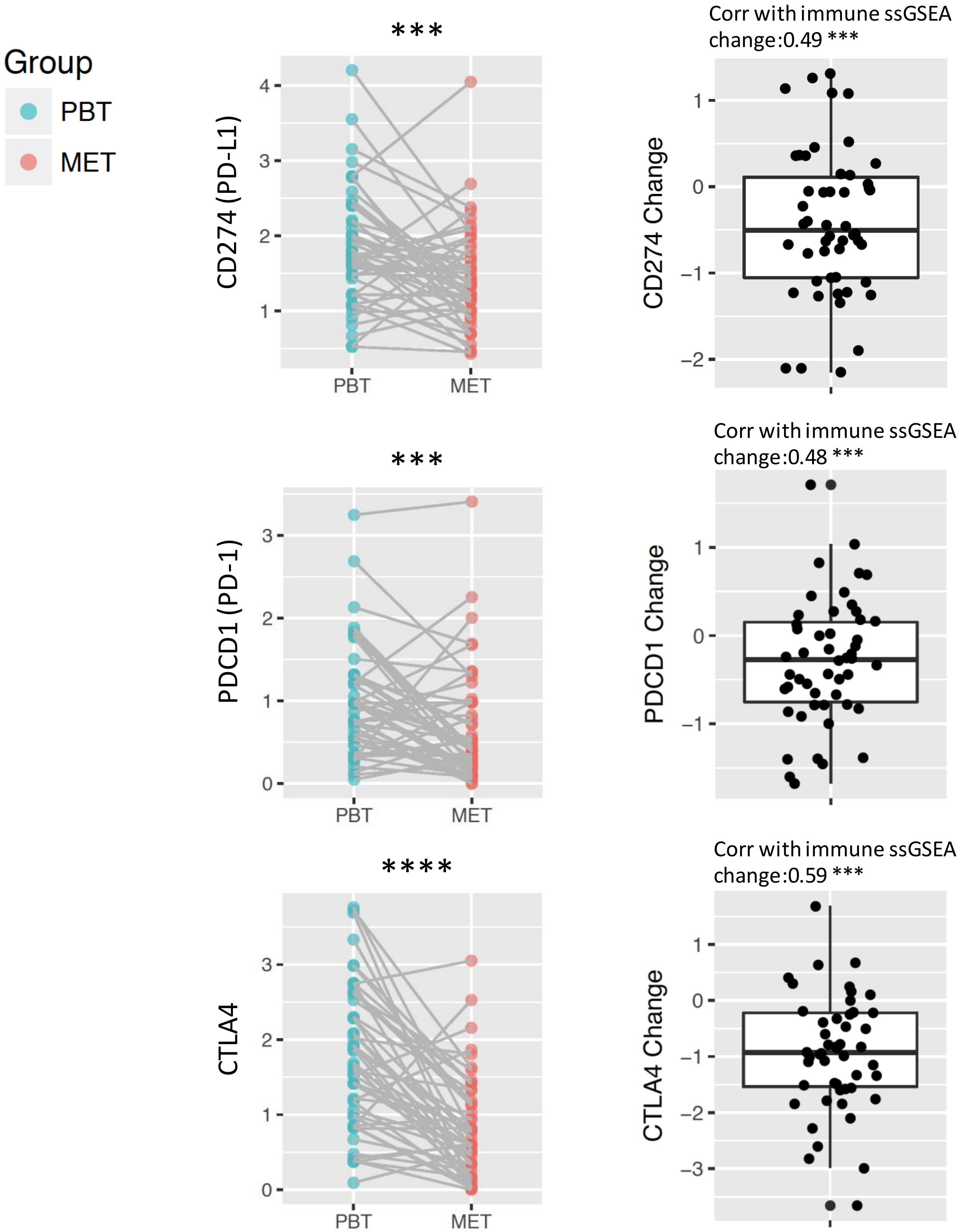


Figure S2. Expressions (log2(TPM+1)) of CD274 (PD-L1), PDCD1 (PD-1), and CTAL4 in PBT and MET. Two-sided Wilcoxon signed rank test was used to compare PBT and MET. Spearman's correlation with immune ssGSEA change was calculated and tested using correlation test. ****p <0.0001, ***p <0.001, **p < 0.01, *p < 0.05.

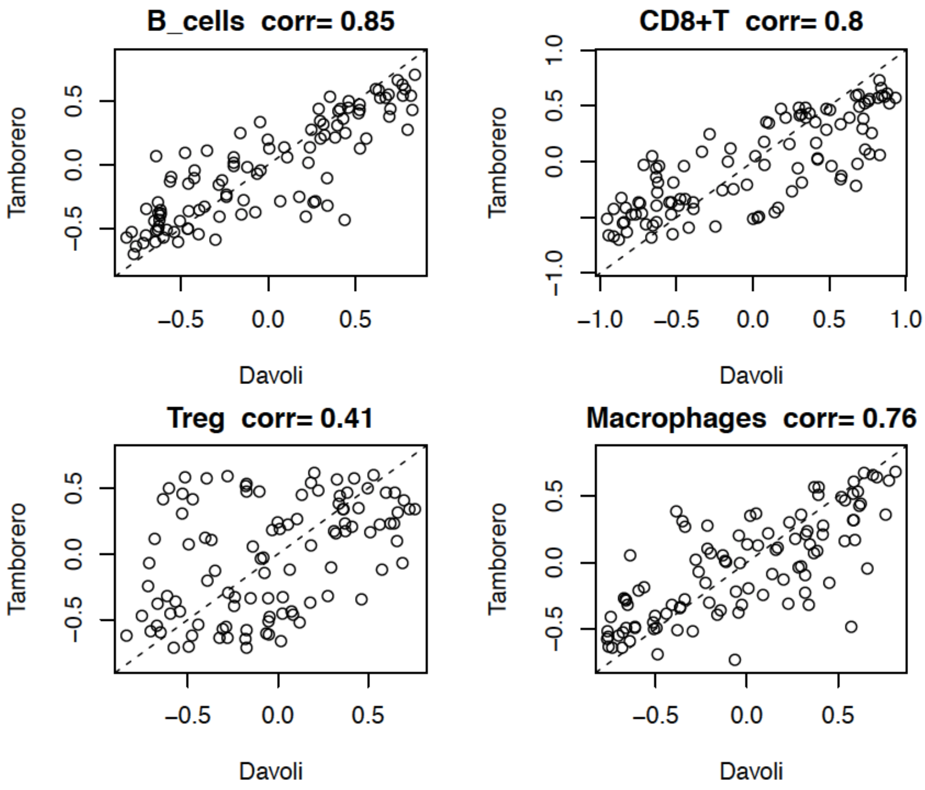


Figure S3. Correlation between GSVA scores of Davoli and Tamborero signatures for PBT/MET pairs in Pan-MET dataset.

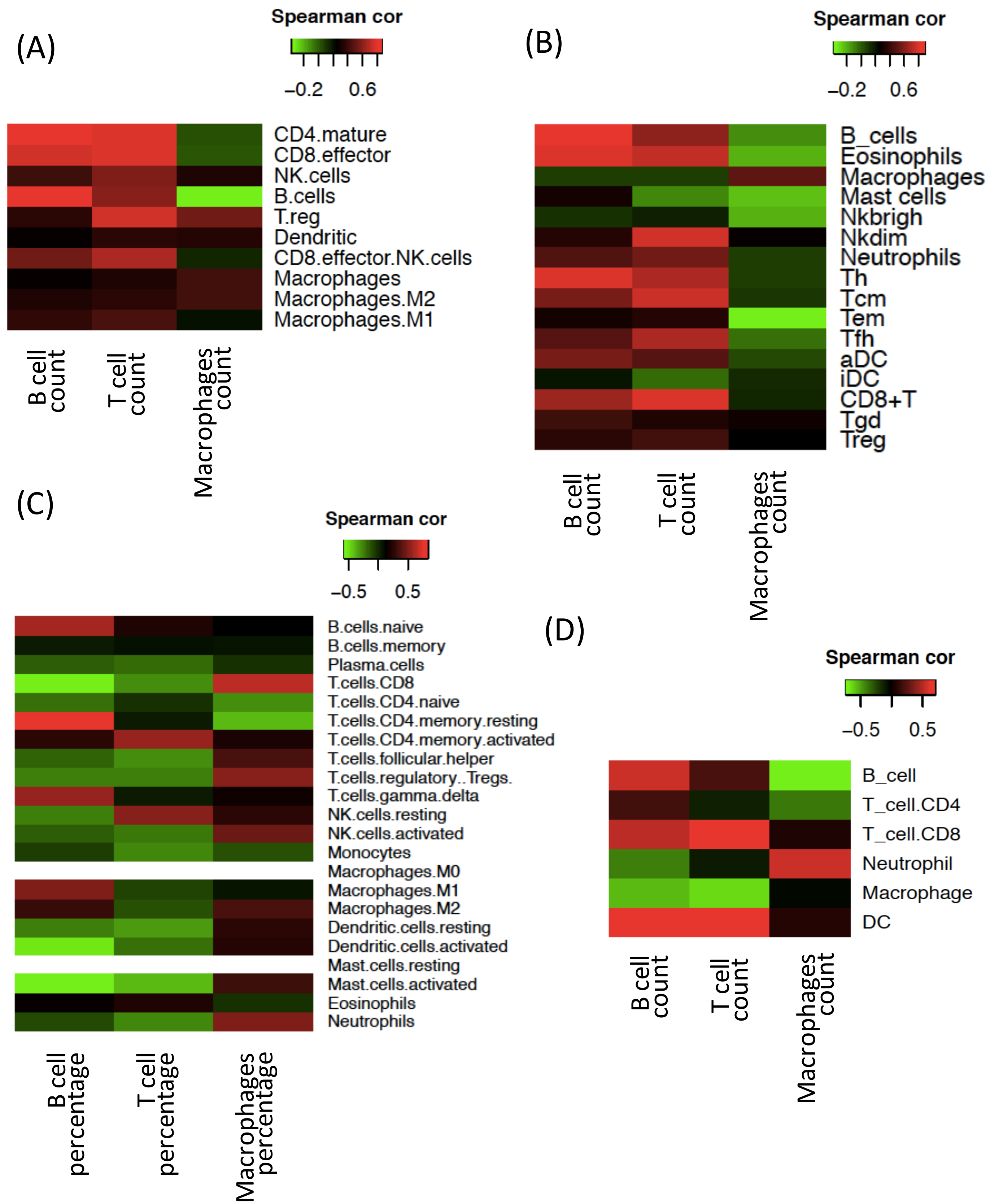


Figure S4. Correlation between immune abundance estimated from RNA-seq data and cell count/proportion (relative to total immune cell count) in single cell RNA-seq dataset. (A-B) GSVA score of (A) Davoli and (B) Tamborero signatures. (C) Percentage relative to total immune level estimated by CIBERSORT. (D) Immune abundance estimated by TIMER. White in the heatmap indicates CIBERSORT estimates are all zero, and spearman's correlation is not applicable.

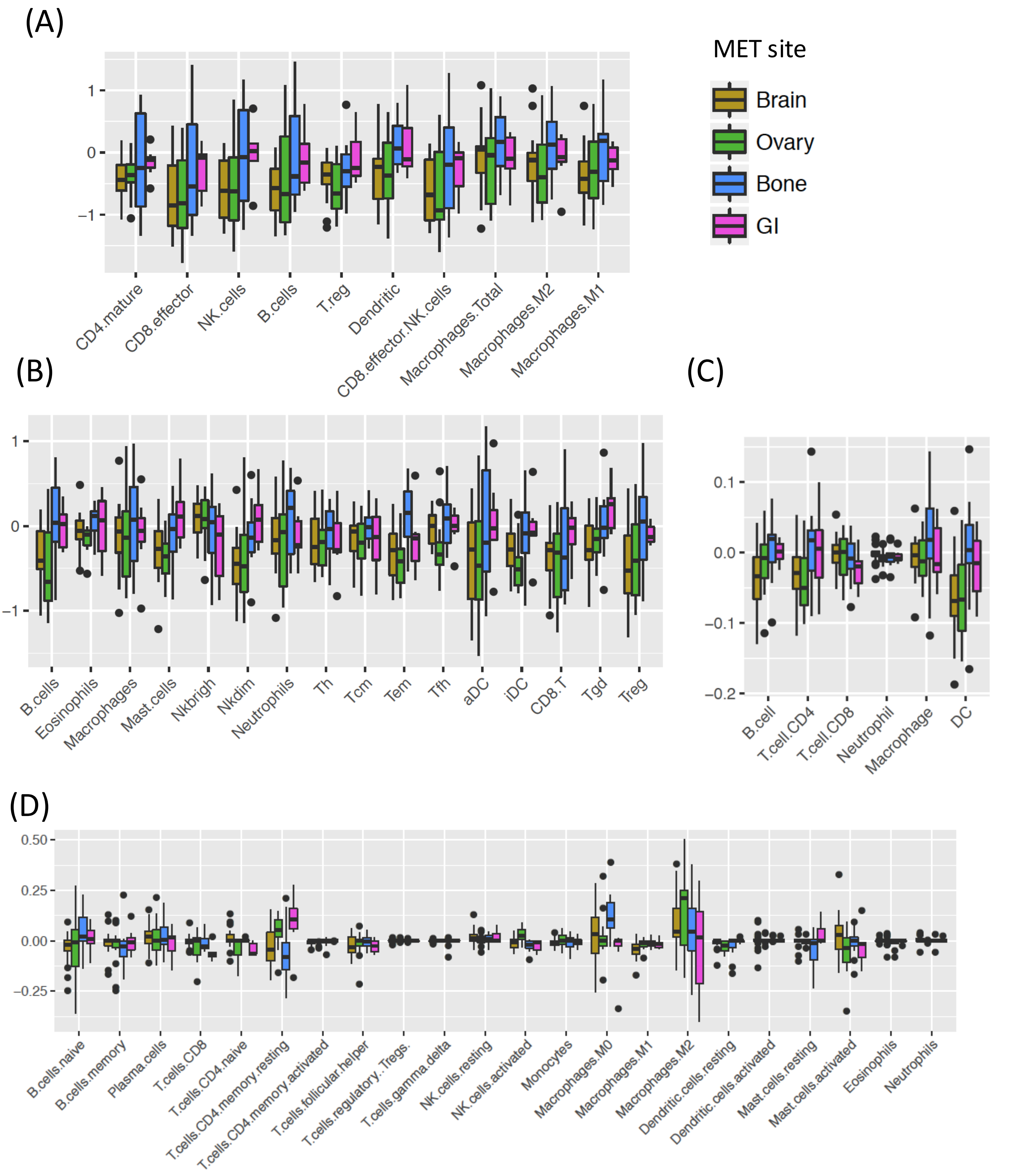


Figure S5. **Comparison of the abundance of immune cell population in PBT/MET pairs in Pan-MET dataset, grouped by MET sites**. (A-B) GSVA score change (MET-PBT) of (A) Davoli and (B) Tamborero signatures. (C) Abundance change estimated by deconvolution method TIMER. (D) Change of percentage relative to total immune estimated by deconvolution method CIBERSORT. ****FDR<0.0001, ***FDR <0.001, **FDR < 0.01, *FDR < 0.05. Two-sided Wilcoxon signed rank test.

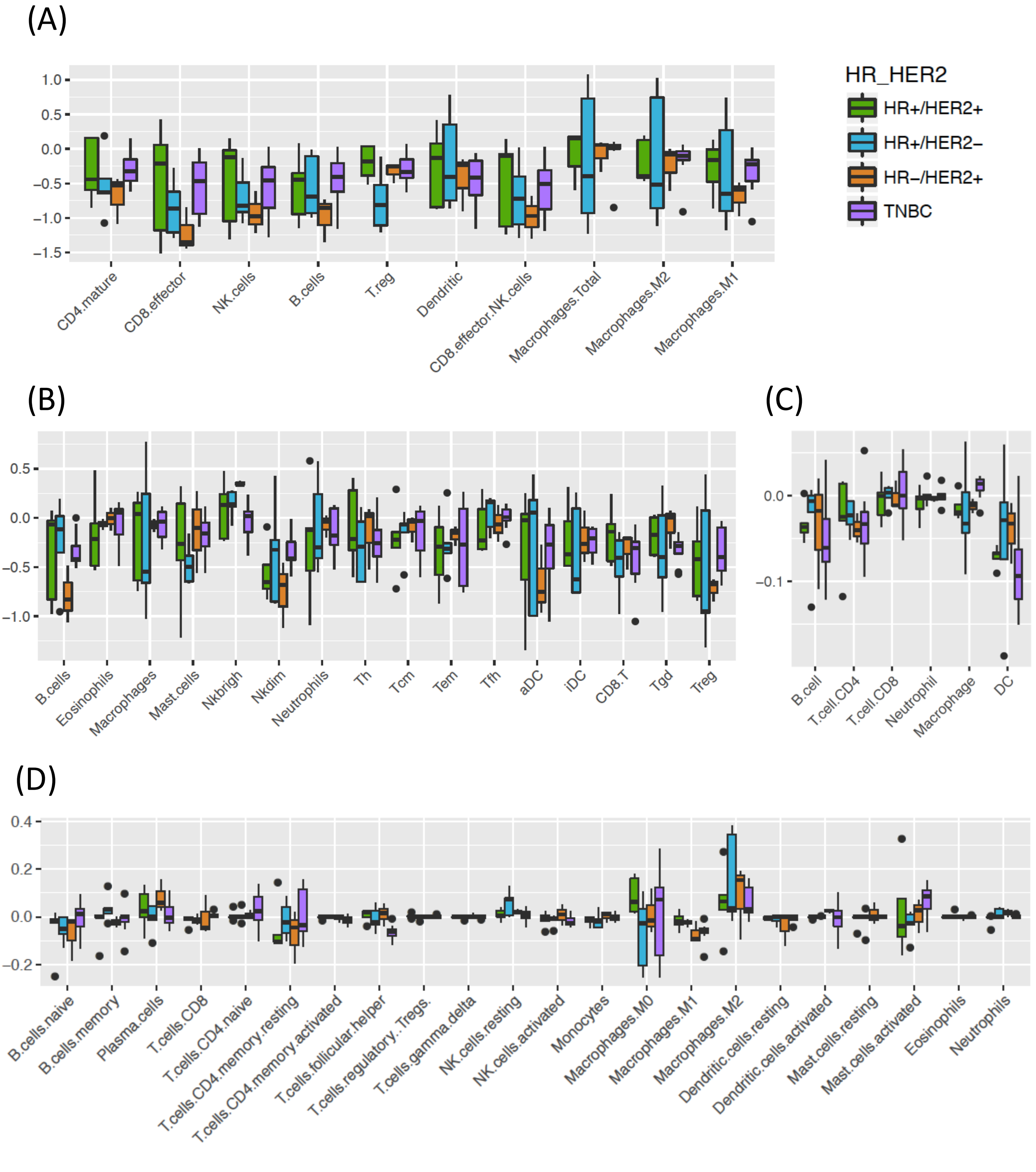


Figure S6. **Comparison of the abundance of immune cell population in PBT/BRM pairs in Pan-MET dataset, grouped by HR/HER2**. (A-B) GSVA score change (BRM-PBT) of (A) Davoli and (B) Tamborero signatures. (C) Abundance change estimated by deconvolution method TIMER. (D) Change of percentage relative to total immune estimated by deconvolution method CIBERSORT. ****FDR<0.0001, ***FDR <0.001, **FDR < 0.01, *FDR < 0.05. Two-sided Wilcoxon signed rank test.

(A)

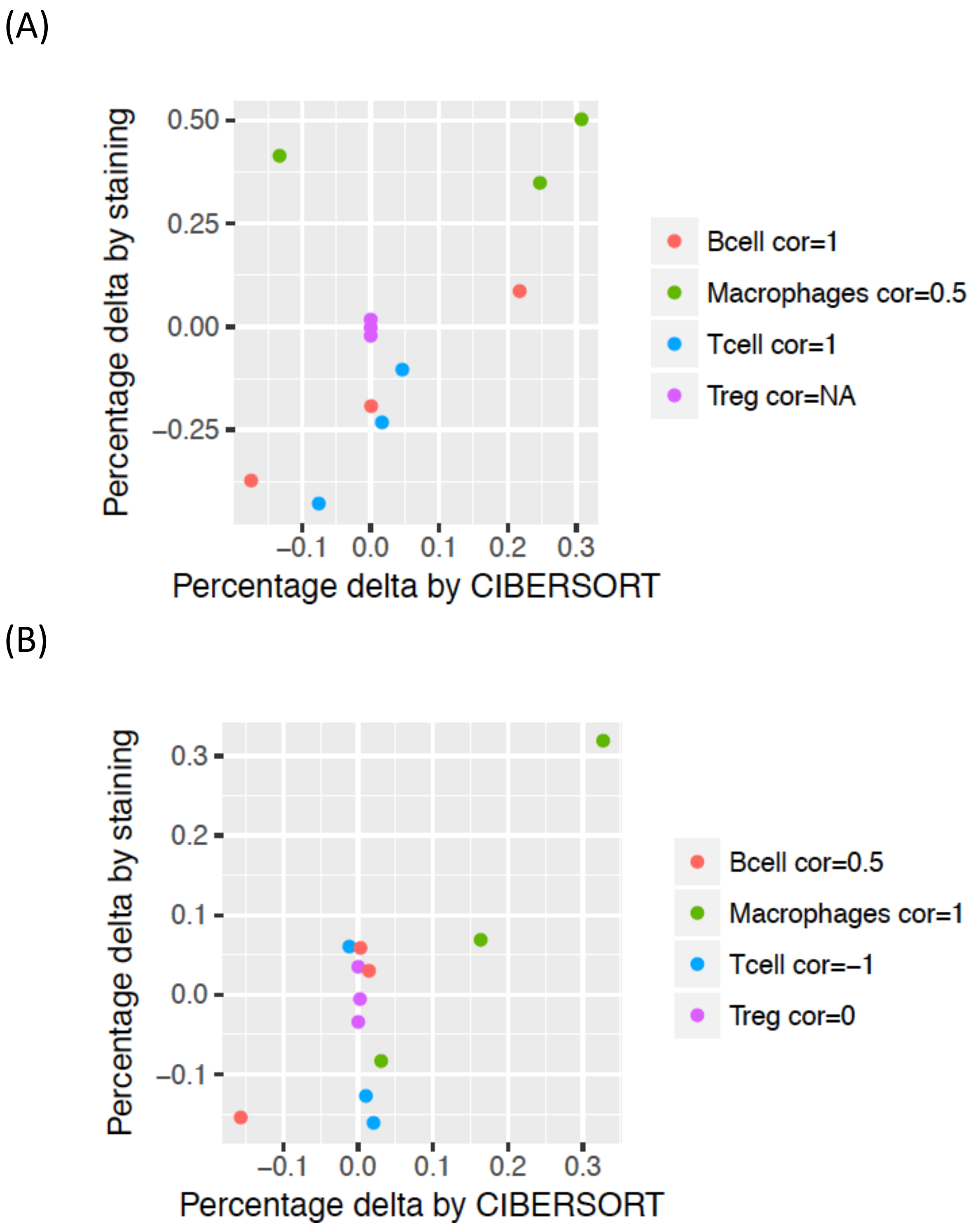


**Figure S7. Correlation between mIHC staining results and CIBERSORT estimates.**
(A) PBT/OVM pairs. (B) PBT/BRM pairs in Pan-MET. Spearman's correlation.

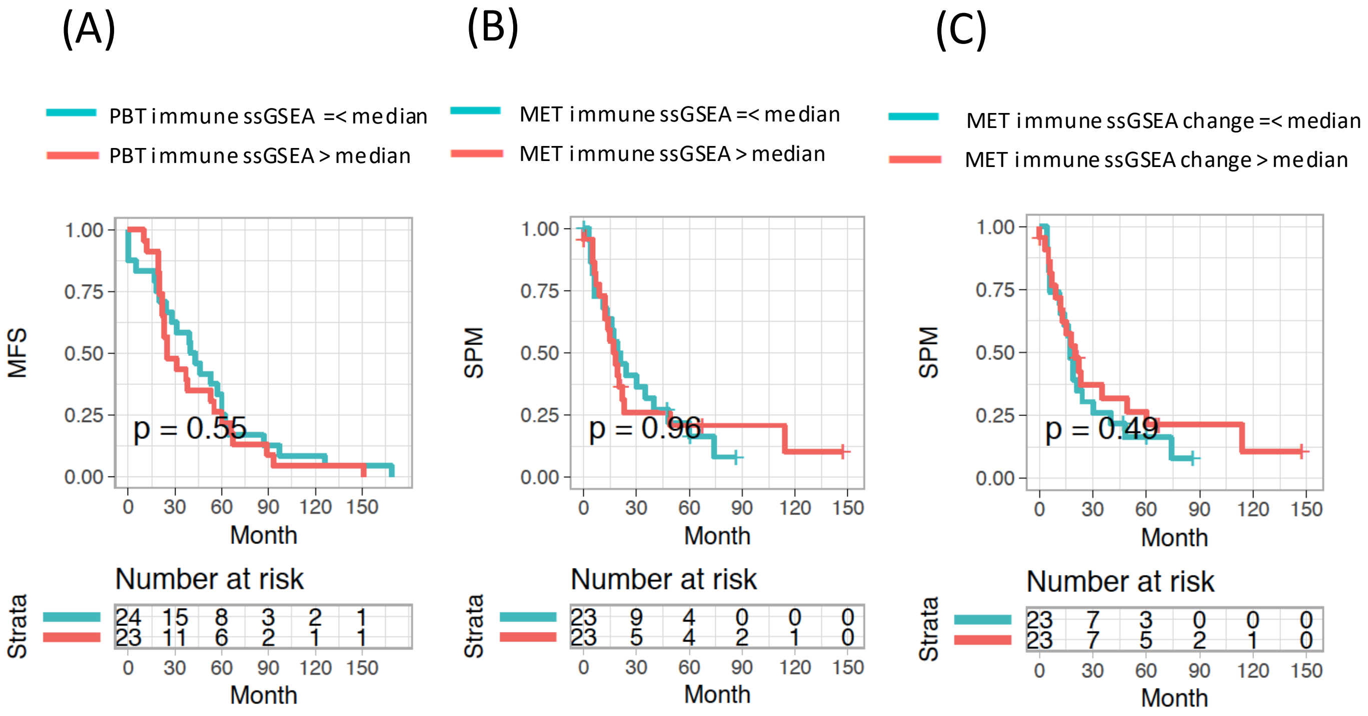


Figure S8. **Test association between survivals and total immune ssGSEA of all pairs of PBTs and METs in Pan-MET dataset.** (A) Kaplan-Meier (KM) curves of MFS for PBTs with total immune ssGSEA below or above median. (B) KM curves of SPM for METs with total immune ssGSEA below or above median. (C) KM curves of SPM for METs with total immune ssGSEA change below or above median. P-values were from log-rank test.

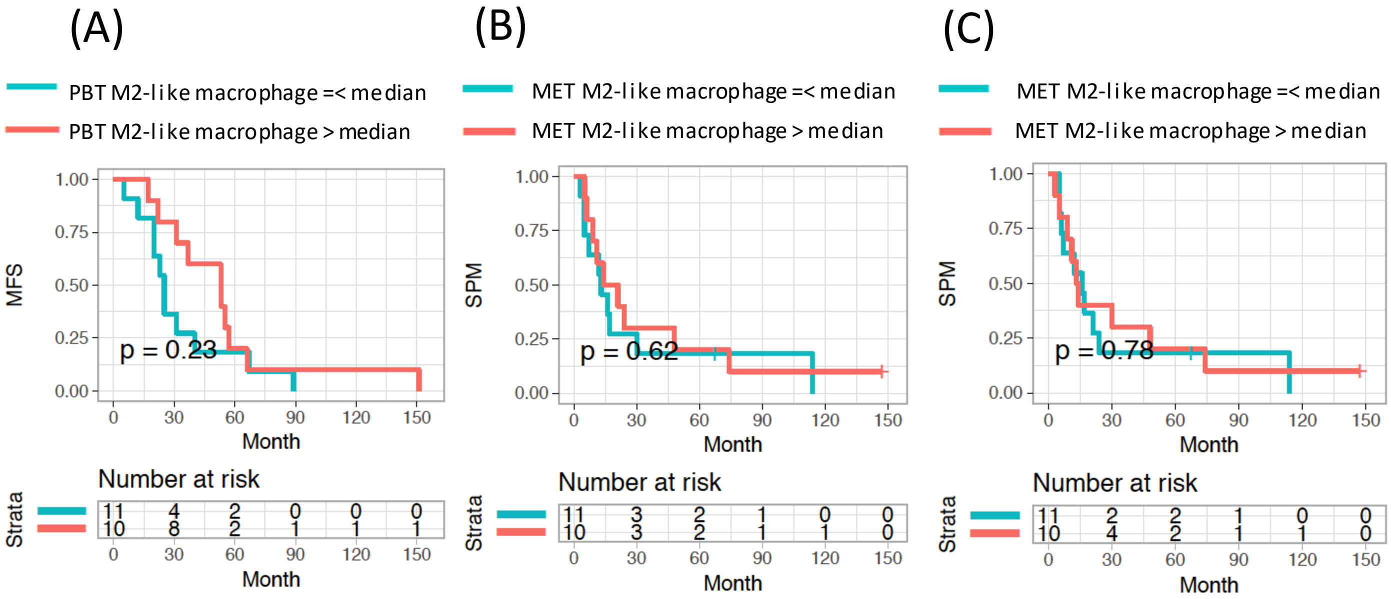


Figure S9. **Test association between survivals and relative percentage of M2-like macrophages of PBT/BRM pairs in Pan-MET dataset.** (A) Kaplan-Meier (KM) curves of MFS for PBTs with relative percentage of M2-like macrophage below or above median. (B) KM curves of SPM for METs with relative percentage of M2-like macrophage below or above median. (C) KM curves of SPM for METs with relative percentage change of M2-like macrophage below or above median. P-values were from log-rank test.
